# Functional gradients dissociate between cortical layers

**DOI:** 10.64898/2026.02.06.704362

**Authors:** J. Karolis Degutis, Jenifer Miehlbradt, Manon Durand-Ruel, Petra S. Hüppi, Dimitri Van De Ville

## Abstract

Functional cortical gradients capture the brain’s large-scale organization along sensory-to-association (G1) and visual-to-somatomotor (G2) axes, yet the laminar circuitry that supports these axes remains largely unknown. Leveraging whole-brain submillimeter 7T resting-state fMRI, we estimated functional connectivity within deep, middle, and superficial cortical depths, integrated these into a multilayer network, and derived depth-resolved functional gradients across the cortex. We then computed an inter-regional dissimilarity index quantifying how distinct each region is from the rest of the cortex, and discovered a systematic dissociation across depth: the superficial-layer index closely followed G1, whereas the deep-layer index aligned with G2. The deep-layer index peaked in receive-dominant regions, consistent with deep layers as principal targets of feedback projections, while the superficial-layer index was maximal in cytoarchitecturally less differentiated transmodal cortex, consistent with its dense recurrent circuitry. Together, these findings demonstrate that functional gradients relate to laminar connectivity, and establish a mesoscale framework for decomposing whole-brain cortical connectivity.

---

Macroscale gradients have become a central framework for characterizing the spatial organization of the human cortex ^1,2^. Resting-state functional connectivity can be described along two principal axes: a sensory-to-association gradient (G1) and a visual-to-somatomotor gradient (G2). These axes align with maps of cytoarchitecture, myelination, receptor density, and developmental change ^3–6^, a convergence that has positioned gradients as a unifying framework for brain organization. However, these gradients capture organization only at a broad spatial scale. Because they are derived from connectivity averaged across the full cortical depth, it remains unknown whether functional macroscale gradients are preferentially expressed within distinct cortical layers.

The neocortex is organized into layers of cortical gray matter that differ and have distinct connectivity patterns and functional roles. The canonical microcircuit characterizes connectivity between areas as feedforward (FF), feedback (FB) or recurrent; FF signals predominantly target middle layers, whereas FB signals project to superficial and deep layers ^7,8^. Laminar measurements in humans have been made possible by ultra-high-field fMRI. Submillimeter acquisitions at 7T have resolved layer-dependent signals in visual ^9^, higher-order association cortices ^10,11^, and more recently across the whole cortex ^12^.

Complementary lines of research have made substantial progress in mapping gradients of laminar organization, including microstructure ^13^, cortical morphology and cytoarchitecture ^3,14^ across the cortex, as well as in resting-state networks ^15,16^. A cortical gradient capturing systematic variation in laminar thickness and laminar neuronal density has been shown to track functional cortical connectivity at rest ^3^. Similarly, the density of excitatory neurons in the superficial layers increases toward association cortices ^6^. These findings suggest that macroscale functional gradients may arise from different cortical layers, yet they have only been linked indirectly, rather than by resolving the functional connectivity across cortical depth.

Here, we test this prediction directly using whole-brain submillimeter 7T resting-state fMRI in 21 healthy adults. For each subject we computed parcel-wise functional connectivity within each of three cortical depths, assembled the matrices into a single multilayer adjacency matrix, and applied joint diffusion-map embedding to obtain functional cortical gradients, each spanning all layers and parcels (Fig. 1A). To quantify the distinctiveness of each parcel’s connectivity profile from the rest of the cortex, we computed an inter-regional dissimilarity index ^18,19^ (Fig. 1B). High values of this index indicate that a parcel’s connectivity differs from the rest of the cortex, while low values indicate that the connectivity patterns are similar. The index was computed either across all layers (Fig. 1B) or within each layer separately (Fig. 1C); the latter case shows the uniqueness of a given area’s layer-specific connectivity.

**Figure 1.**
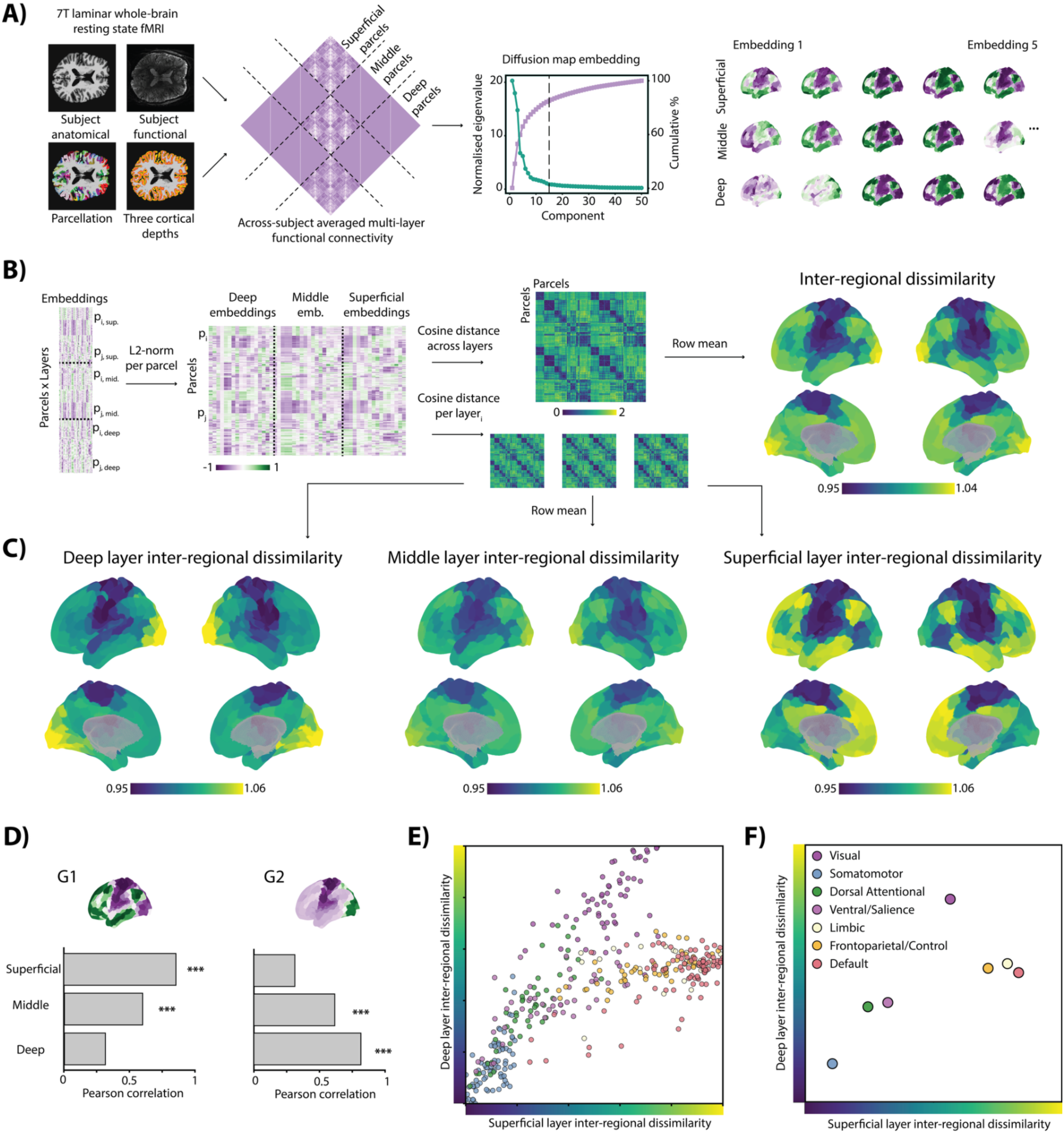
Layer-specific dissimilarity indices. **A)** Processing pipeline. Anatomical image, resting-state functional images, Schaefer-400 parcellation, and a cortical depth map computed (all in subject functional space) were used to extract parcel time series from three equivolume depth bins (deep, middle, superficial layers). For each depth, parcel-wise Pearson correlations were computed to build a multilayer adjacency matrix per subject, which were then averaged across subjects. Joint diffusion map embedding using a cosine-kernel was applied. Fifteen low-dimensional embeddings (five shown) across layers were estimated. **B)** Inter-regional dissimilarity. Cosine distances were computed between parcels using embeddings of all layers. The dissimilarity matrix was averaged per row. **C)** Layer-specific inter-regional dissimilarity. Computed as in B), but for each layer’s embeddings individually. **D)** Correlation between layer-specific inter-regional dissimilarity indices and the canonical sensory-association axis (G1) and visual-somatomotor axis (G2). **E)** Relationship between superficial and deep inter-regional indices across parcels. Each point is a parcel colored by its resting-state network using the seven network Yeo parcellation ^17^ **F)** Resting-state network centroids from E). *** indicate pspin<0.0002.

The across-layer index was high in visual cortex, intermediate in association cortex, and low in somatomotor cortex, resembling a combination of the two principal functional connectivity gradients (Fig. 1B). When computed per layer, the two gradients dissociated: the superficial-layer index corresponded closely to the canonical sensory-to-association axis, with low values in sensory and motor regions and high values in association cortex, while the deep-layer index corresponded to the visual-to-somatomotor axis (Fig. 1C). We quantified how similar the layer-specific indices are by correlating them with G1 and G2 (Fig. 1D): the superficial-layer index correlated with G1 (r=0.86, pspin<0.0002), but not G2 (r=0.31, pspin=0.11); the deep index showed the opposite pattern (G1: r=0.32, pspin=0.054; G2: r=0.82, pspin<0.0002), while the middle-layer index correlated with both gradients, but more weakly (G1: r=0.60, pspin<0.0002; G2: r=0.62, pspin<0.0002). The superficial and deep indices had the weakest correlation (r=0.67, pspin=0.001), while the middle correlated with both the superficial (r=0.89, pspin<0.0002) and the deep index (r=0.90, pspin<0.0002).

Given the greatest divergence between superficial and deep maps, we compared them directly. The two indices separated parcels by their Yeo-7 resting-state network membership ^17^ (Fig. 1E–F; F = 53.7, pspin<0.0002). Superficial dissimilarity was highest in control, limbic, and default networks; deep dissimilarity peaked in visual cortex (Fig. 1E–F). Layer effects were significant within every network (Fig. 2A), and the dissociation was preserved in all but one subnetwork using the Yeo-17 solution ^17^ (Supplementary Fig. 1).

**Figure 2.**
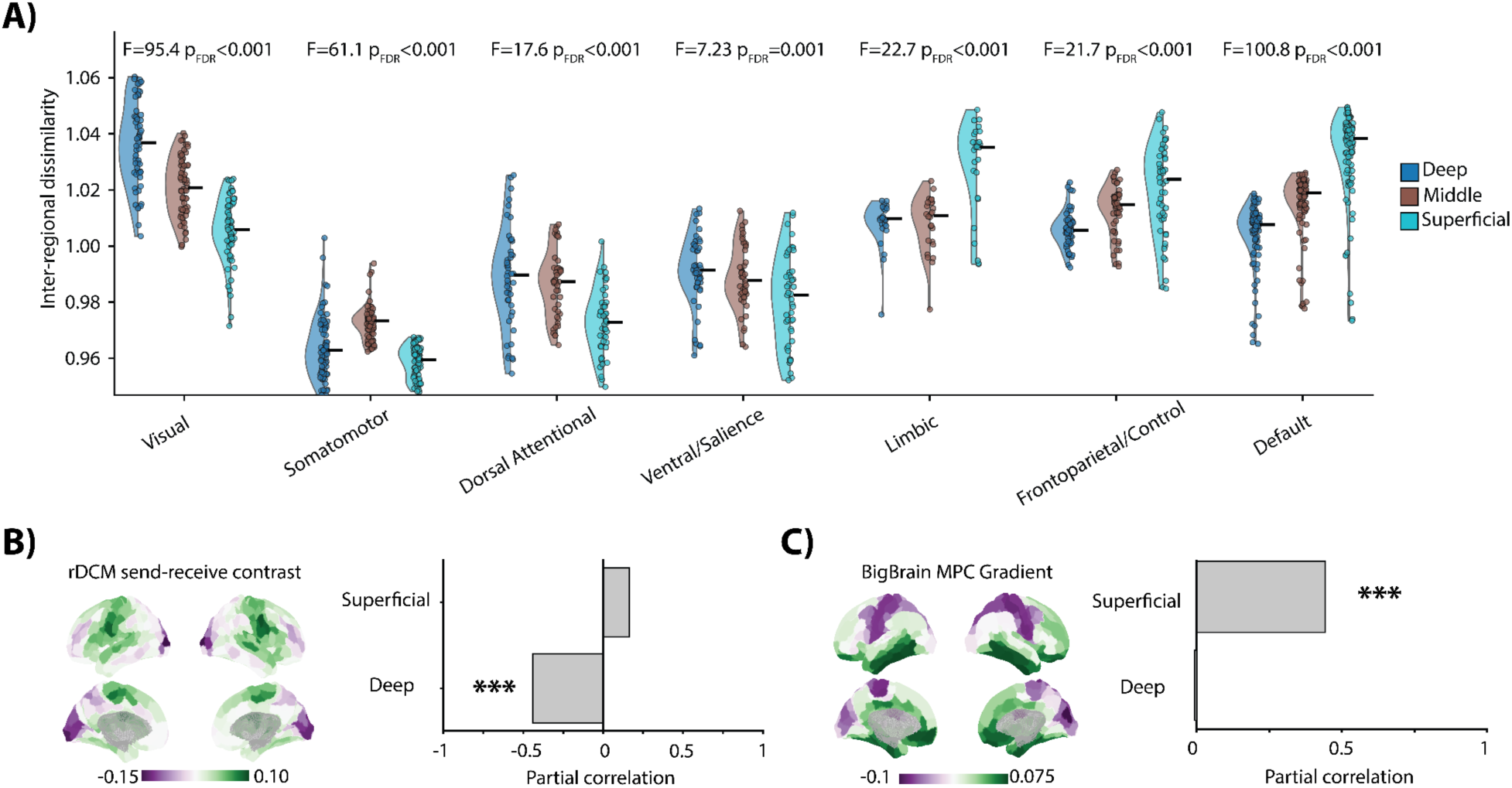
Layer-specific index network, hierarchy, and structural relations. **A)** Layer-specific inter-regional dissimilarity indices mapped per network. Each point represents a parcel within a given network. Statistics show tests for layer effects per network using a one-way ANOVA (FDR-corrected across networks). **B)** Left: cortical map of a send-receive contrast derived based on regression dynamic causal modeling (rDCM). Right: partial correlation of layer-specific inter-regional indices and rDCM send-receive contrast when controlling for the other layer-specific index. **C)** Left: cortical map of the BigBrain microstructural profile covariance (MPC) gradient. Right: partial correlation of layer-specific inter-regional indices and MPC gradient when controlling for the other layer-specific index. *** indicate pspin<0.0002.

Next, we asked whether the layer-specific indices reflect a signal-propagation hierarchy. Using a previously published effective-connectivity send-receive asymmetry map ^3^, in which negative values mark receive-dominant regions, we computed partial correlations between each layer’s index and the asymmetry, controlling for the other layer. Only the deep-layer index showed a significant negative association (superficial: *r* = 0.057, pspin=0.36; deep: *r* = -0.57, pspin<0.0002; Fig. 2B): regions with high deep-layer dissimilarity were those that are receive-dominant at rest. This is consistent with the canonical microcircuit, in which deep layers are recipients of feedback projections.

We then asked whether the indices track cortical microstructure, using the BigBrain microstructural profile covariance (MPC) gradient ^3,15^, which captures a sensory-to-limbic transition in cytoarchitectural differentiation. High values in this index indicate areas that are less cytoarchitectonically differentiated. Controlling for the other layer, only the superficial-layer index correlated positively with the MPC gradient (superficial: r = 0.45, pspin=0.0041; deep: r = -0.0044, pspin=0.51; Fig. 2C), suggesting that less differentiated association areas have unique interareal superficial-layer connectivity.

Several analyses support the robustness of these results. Temporal signal-to-noise (SNR) ratio decreased from deep to superficial layers (Supplementary Fig. 2A, C–D), but did not correlate with the corresponding layer’s dissimilarity index (Supplementary Fig. 2B), indicating that the laminar dissociation is not driven by SNR. Varying the number of gradient components used to construct the indices yielded highly similar maps from eight gradients onwards (Supplementary Fig. 3A) and maintained the double-dissociation between superficial and deep indices when correlated to G1 and G2 (Supplementary Fig. 3B). In random split-half resamplings of the 21 subjects, each layer-specific index correlated highly with its split counterpart (Supplementary Fig. 4). The dissociation depended on coupling each region to itself across depths in the multilayer matrix: it was clearest at high coupling values and the dissociation reduced when the coupling was low (Supplementary Fig. 5). We also computed the dissimilarity indices on HCP MMP 1.0 parcellation and find similar layer-specific maps (Supplementary Figure 6); average pairwise distances between resting-state network centroids of deep and superficial indices in the HCP MMP 1.0 parcellation are similar to the Schaefer-400 parcellation centroid distances (r=0.93, pspin=0.0003).

Macroscale cortical organisation has increasingly been described in terms of convergence; i.e., functional, cytoarchitectonic and anatomical hierarchies align along a single sensory-to-association axis ^20^. Our findings provide a more nuanced perspective on this view. When resting-state connectivity differences are resolved across cortical depth, the two principal gradients dissociate: the sensory-to-association axis is preferentially present in superficial layers, whereas the visual-to-somatomotor axis is predominant in deep layers. Macroscale functional gradients thus appear to arise from laminar connectivity and become spatially indistinguishable when the signal is averaged across cortical depth.

The association between superficial layers and G1 may reflect the recurrent circuitry of transmodal cortex, where dense local interconnection allows activity to be sustained and circulated within networks. Superficial layers appear particularly important for these dynamics: in non-human primates, gamma-band activity in superficial lateral prefrontal cortex carries working-memory information, indicating a role in maintaining internally generated representations ^21,22^. Anatomical work in mice extends the link to the default mode network, where layer 2/3 neurons in DMN regions project to other DMN regions ^23^; because the default network occupies the same end of the superficial axis, similar recurrent loops may support self-generated cognition ^16^. The MPC correlation reinforces this interpretation as granular differentiation diminishes towards limbic and association cortex, and therefore the superficial-layer index becomes increasingly prominent.

The selective coupling between deep-layer dissimilarity and the effective-connectivity send-receive contrast supports a complementary interpretation for G2. Regions with high deep-layer distinctiveness are receive-dominant at rest, consistent with deep layers as canonical targets of feedback projections ^7–9^. Since this signature is strongest in visual cortex at rest, it may reflect ongoing top-down modulation of unimodal sensory areas. Such a pattern would be expected if intrinsic activity in early sensory cortex is continually shaped by descending signals from higher areas, which is consistent with hierarchical accounts in which deep-layer activity carries contextual or predictive influences from downstream regions ^9,24^.

We measured the laminar functional signal using gradient-echo BOLD, which is biased toward superficial layers as it carries signal that drains from deeper layers toward the pial surface; this limits the anatomical specificity of the superficial depth ^25^. However, this bias would actually predict more similar responses between deep and superficial layers: by transporting deep-layer signal into the superficial depths, venous drainage should make the two profiles more alike and blur depth-dependent maps. The dissociation we observe is therefore opposite to the prediction based on venous contamination, and is likely a conservative estimate of the true laminar separation. This dissociation also persisted across our robustness analyses, suggesting that laminar decomposition provides a principled mesoscale description of macroscale cortical gradients.

Overall, the present work demonstrates that laminar indices derived from multilayer functional gradients offer an informative characterization of how layer-resolved functional architecture varies across intrinsic networks. These findings suggest that laminar functional gradients provide a valuable framework for studying how cortical layers contribute to large-scale communication at rest and for future investigations of how hierarchical laminar organization may differ across individuals, developmental stages, and clinical conditions.

## Methods

### Participants

We scanned 21 healthy young participants (18 to 22 years, 11 female). All participants gave written informed consent and received monetary compensation for their participation. The study was approved by the ethics committee of the Canton of Geneva (CCER). Ethical regulations relevant to human research participants were followed.

### Acquisition

MRI data were acquired on a Siemens MAGNETOM Terra.X 7T MRI system (Siemens, Erlangen, Germany) using an 8Tx/32Rx head coil (Nova Medical) for clinical exams. T1-weighted anatomical images were obtained with MP2RAGE (TR=5000 ms, TE=2.27 ms, 0.75 mm isotropic; TI1/2=900/2700 ms; flip angles=3°/5°) ^26^. Following ^27^, high-resolution resting-state fMRI was acquired with single-shot T2*-weighted 2D gradient-echo EPI at 0.8 mm isotropic resolution (FOV=192 mm; base matrix=240; 135 interleaved slices with 10% gap; AP phase-encode; receiver bandwidth=1226 Hz/Px; echo spacing=1.00 ms; EPI factor=240; partial Fourier=6/8). Parallel imaging used in-plane acceleration R=3 together with simultaneous multi-slice of 3. The resting-state run had a TR=3290 ms, TE=29 ms, flip angle=80°, 138 volumes resulting in 7.6 minutes of acquisition per subject. Subjects 17 and 18 had fewer acquired volumes: 114 and 131, respectively.

### Anatomical preprocessing

We processed the data using a previously developed automated laminar fMRI preprocessing pipeline ^10,28^. To obtain accurate reconstructions of gray and white matter surfaces from structural MP2RAGE images, we applied a Freesurfer-based workflow. First, a bias-corrected version of the INV2 image was multiplied with the UNI image to create a T1-weighted image that was both spatially homogeneous and free of extracerebral noise (MPRAGEize ^29^). This T1-weighted image was then passed to CAT12 for tissue segmentation ^30^. A high-quality brain mask was generated by combining the segmented gray and white matter compartments. Subsequently, the T1-weighted image was processed using Freesurfer’s recon-all pipeline, with the -hires flag enabled to account for the high spatial resolution ^31^. We substituted Freesurfer’s automatically generated brain mask with the more accurate CAT12-derived brain mask.

For parcellation, each participant’s segmented structural surfaces were transformed into fsLR164k CIFTI space using ciftify recon-all, part of the Python-based ciftify package ^32^. Expert settings were used to enable registration to the high-resolution 0.5 mm MNI152 T1w template. Ciftify adapts the HCP minimal preprocessing pipeline to non-HCP datasets and includes surface-based alignment via MSMSulc to HCP’s fs_LR surface space.

To align structural and functional data, we calculated the transformation between the Freesurfer-processed T1-weighted image and the mean volume of the GE-BOLD run using ANTs ^33^. This registration consisted of an initial affine transformation followed by a nonlinear warping step using the Symmetric Normalization (SyN) algorithm, which also served for distortion correction. Laminar segmentation was performed using LAYNII ^34^. We computed three equivolume depth bins across the cortical ribbon in functional space.

### Functional preprocessing

For 12 subjects for whom a phase image was acquired, we denoised each resting-state run before motion correction with NORDIC PCA (MATLAB NIFTI_NORDIC, default parameters), using the magnitude EPI time series together with its corresponding phase series. NORDIC applies locally low-rank PCA with distribution-corrected thresholds to selectively suppress thermal noise while preserving spatial precision and thus improving tSNR ^35^. This has been validated for sub-millimeter 7T fMRI ^36^. We also found improvement in tSNR for subjects with NORDIC (Supplementary Figure 2C-D).

Resting-state functional scans were motion-corrected using AFNI’s 3dVolreg, employing Fourier interpolation ^37^. Volumes were aligned to the timepoint with the fewest outlier voxels, as identified by 3dToutcount, following standard AFNI preprocessing protocols. Preprocessing of the resting-state fMRI time series was performed in SPM12 ^38^. Nuisance regressors included a constant term, linear and quadratic trends, mean white-matter signal, mean CSF signal, rigid-body motion parameters, and their first temporal derivatives. A high-pass filter of 0.01 Hz was applied. Model estimation was run per subject. The residual images were used for subsequent resting-state analyses.

### Atlases

Schaefer-400 ^39^ and HCP MMP 1.0 atlas ^40^ labels were transformed from fs_LR to each subject’s native freesurfer surface space using HCP’s wb_command ^41^ and to the subject-specific surface generated by ciftify’s recon-all function. These were then projected into each subject’s functional volume space using HCP workbench relying on the subject-specific surfaces in functional space.

### Laminar resting-state functional connectivity matrix

To examine the spatial structure of laminar functional connectivity, we built a multilayer functional connectivity matrix by computing Fisher z-transformed correlation coefficients for each subject, averaging these values across subjects, and then applying the inverse Fisher transform. Each layer-specific adjacency matrix was positioned along the diagonal, while the off-diagonal blocks had identity matrices with a set weight of one, thus forming a connection across layers within each corresponding region. Using BrainSpace ^42^, a cosine-kernel derived affinity matrix was calculated from the multilayer adjacency matrix. A sparsity threshold was applied to retain the top 10% of strongest connections per row. Diffusion map embedding was then performed on the resulting affinity matrix to extract cortical gradients, the principal dimensions of variance in laminar connectivity patterns across cortical regions and layers, that captured 85% of the cumulative eigenvalue spectrum. These gradients provide a low-dimensional representation of the spatial connectivity landscape for each layer. Off-diagonal block identity matrices enforced alignment of the same region across depths so gradients would reflect laminar rather than global differences (Supplementary Figure 5).

### Inter-regional dissimilarity index

Following ^19^, we computed an inter-regional dissimilarity index to quantify how distinct each parcel’s laminar connectivity profile is from the rest of the cortex. The joint diffusion-map embedding provides, for every parcel, gradient coordinates at each cortical depth. Within each layer, gradient components were z-scored across parcels before L2-normalization. This centers each component and places all retained gradients on a common scale, so that the dissimilarity index reflects the full multivariate embedding rather than being dominated by the leading, highest-variance gradient. Standardizing each layer separately keeps this weighting consistent across depths, so that a layer’s contribution to the laminar profile is governed by its spatial connectivity pattern and not by between-layer differences in embedding scale.

The layer-specific index was computed from a single layer’s embedding: pairwise cosine distances were calculated between all parcels’ unit-normalized layer vectors, yielding one cosine-distance matrix per layer (deep, middle, superficial). A parcel’s layer-specific inter-regional dissimilarity was defined as the mean cosine distance between its layer profile and those of all other parcels; the mean of its row in that layer’s matrix (Figure 1C). High values indicate that a parcel’s connectivity within that layer is distinct from the rest of the cortex.

The across-layer index was computed analogously, but each parcel’s laminar profile was formed by concatenating its unit-normalized vectors from all layers; dissimilarity was the mean, across all other parcels (Figure 1B).

### Macroscale functional gradients, effective connectivity map and BigBrain gradient

The two principal macroscale connectivity gradients were taken from the canonical resting-state functional connectivity embedding of Margulies et al. ^1^ to provide reference axes independent of our laminar embeddings. They were retrieved in fsLR space at 32k density using the neuromaps toolbox ^43^. Both maps were parcellated to the Schaefer-400 atlas ^39^ with the neuromaps *Parcellater*, after converting the atlas to fsLR-space surfaces.

We used an effective connectivity matrix from previously published studies ^3,15^ that used data from the Microstructure-Informed Connectomics (MICs) cohort ^44^, as well as from a replication sample based on the Human Connectome Project dataset ^45^. Effective connectivity among the Schaefer-400 cortical parcels was estimated from resting-state fMRI data using regression dynamic causal modelling (rDCM) ^46^. Following ^3^, the send-receive effective connectivity matrices were used to quantify asymmetry-based hierarchical organization. The asymmetry hierarchy was computed as the difference between each region’s receive in-degree and send out-degree, with higher positive values indicating parcels that preferentially send, while negative values indicate parcels that receive.

We used the histology-derived BigBrain microstructural profile covariance (MPC) gradient ^4^. The authors of the original study sampled intensity profiles along 50 equivolumetric intracortical surfaces of the 100 µm BigBrain reconstruction ^47^, and an MPC matrix was built by correlating depth-wise intensity profiles while controlling for the cortex-wide mean profile. The MPC was thresholded row-wise to the top 10% of entries, converted to a normalized-angle affinity matrix, and embedded with diffusion maps to obtain the principal gradient that captured a sensory-to-limbic axis of cytoarchitectural differentiation. We obtained the microstructural gradient for the Schaefer-400 parcellation using the ENIGMA Toolbox ^48^.

### Robustness analyses

To characterize potential influences of layer-dependent signal quality, we computed temporal SNR (tSNR) across the three depth bins and repeated this for the full sample and for subgroups with and without NORDIC processing. We computed group-average parcel tSNR maps for each layer, per-subject mean values for each layer (averaged across parcels), and a one-way ANOVA across layers on these per-subject layer means (Supplementary Figure 2).

We tested whether laminar dissimilarity indices computed on the Schaefer-400 atlas were sensitive to the number of gradients used to construct them by repeating the full gradient workflow while varying the number of gradient components from 5 to 25. We re-estimated all indices and computed a correlation matrix per index across gradients, and additionally correlated the deep and superficial indices with G1 and G2 at each gradient (Supplementary Figure 3).

We assessed the reproducibility of the laminar dissimilarity indices by repeatedly splitting the sample into independent halves. Across 500 iterations, subjects were randomly partitioned into two groups, a group-average functional connectivity matrix was constructed for each half, and the full gradient workflow was run separately on each (retaining the original number of gradient components). We then recomputed all dissimilarity indices and quantified their agreement between halves as the absolute Pearson correlation, yielding a distribution of split-half reliabilities for each index, which we summarised as raincloud plots (Supplementary Figure 4).

We tested whether the laminar dissimilarity indices were sensitive to the inter-layer coupling weight used to assemble the multilayer adjacency matrix. The off-diagonal blocks of this matrix link each parcel to itself across cortical depths, and in the main analysis this coupling was fixed to a weight of one. We repeated the full gradient workflow while varying this weight from 0 to 1 in steps of 0.1, holding the number of gradient components fixed at the value used in the main analysis. For each weight we rebuilt the multilayer adjacency matrix, re-estimated the cortical gradients, and recomputed all inter-regional dissimilarity indices. Following the gradient robustness analysis, we correlated the deep and superficial indices with G1 and G2 at each weight (Supplementary Figure 5).

To assess whether the relative arrangement of network centroids was consistent across atlases, we compared centroid geometry derived from the Schaefer-400 and HCP MMP 1.0 atlases (Supplementary Figure 6). Within each dataset, we computed all pairwise Euclidean distances between centroids and vectorized the upper triangle of the resulting distance matrix. We correlated the two distance vectors. Statistical significance was evaluated with a label-permutation null in which the centroid labels of the second dataset were randomly permuted and the correlation recomputed.

### Statistical tests

Pairwise relationships between cortical maps were quantified using Pearson correlation computed across Schaefer-400 parcels. Spatial autocorrelation was accounted for using a spin test ^49^ implemented in the ENIGMA Toolbox ^48^: Schaefer-400 parcel centroids on the Conte69 spheres were rotated, generating a permutation index that reassigns parcel values to rotated positions. Spin-based p-values were obtained using ENIGMA’s *perm_sphere_p*, which evaluates the empirical correlation against the null distribution obtained from the rotated parcels. This procedure was used for the following contrasts: pairwise comparisons of layer-specific inter-regional indices (superficial vs. middle, superficial vs. deep, and middle vs. deep), layer-specific inter-regional indices and G1 or G2, and layer-specific inter-regional indices and layer-matched tSNR maps. Each contrast used 5000 rotations with a fixed random seed.

To test whether parcelwise effects differed across resting-state networks as a function of the deep and superficial dissimilarity indices, we fit a two-factor model with network (seven resting-state networks) and index (deep and superficial) and evaluated the network and index interaction using an F-test. Statistical significance was assessed using spatial spin permutations implemented using ENIGMA’s *rotate_parcellation* on Schaefer-400 parcel centroids defined on the Conte69 spherical surfaces. For each of 5000 rotations, the same Schaefer-400 permutation index was applied to all indices to preserve cross-index dependence.

To isolate layer-specific associations with the rDCM send-receive contrast and the BigBrain gradient while accounting for shared variance across the deep and superficial indices, partial correlations were computed for the deep and superficial layer-specific inter-regional index. For each test, covariates (the other layer-specific inter-regional map) were regressed from both the target laminar map and the relevant external map (rDCM contrast or BigBrain gradient); the empirical partial correlation was then defined as the Pearson correlation between the resulting residuals. As in the previous correlations, statistical significance was assessed with the same ENIGMA spin-test approach using 5000 rotations, yielding a spin-based p-value for the residual-residual association.

## Contributions

Conceptualization: J.K.D.; Methodology: J.K.D., D.V.D.V; Formal Analysis: J.K.D.; Software: J.K.D.; Visualization: J.K.D.; Investigation: J.M., M.D-R.; Project Administration: J.M., M.D-R.; Funding Acquisition: P.S.H., D.V.D.V.; Writing - Original Draft Preparation: J.K.D.; Writing – Review & Editing: J.K.D., J.M., P.S.H., D.V.D.V.; Supervision: P.S.H., D.V.D.V.

## Data Availability Statement

Preprocessed data will be made available on the OSF. The analysis code can be found on GitHub (https://github.com/degutis/corticalGradients_layers).

## Acknowledgements

We acknowledge the MRI Platform of the FCBG (Fondation Campus Biotech Geneva) and the CIBM Center for Biomedical Imaging for providing expertise and resources to conduct this study. The study was supported by Swiss National Science Foundation (SNFS) grant no. 212714. We thank Francesca Saviola for extensive discussions and her feedback on the manuscript.

## Supplementary Figures

**Supplementary Figure 1.**
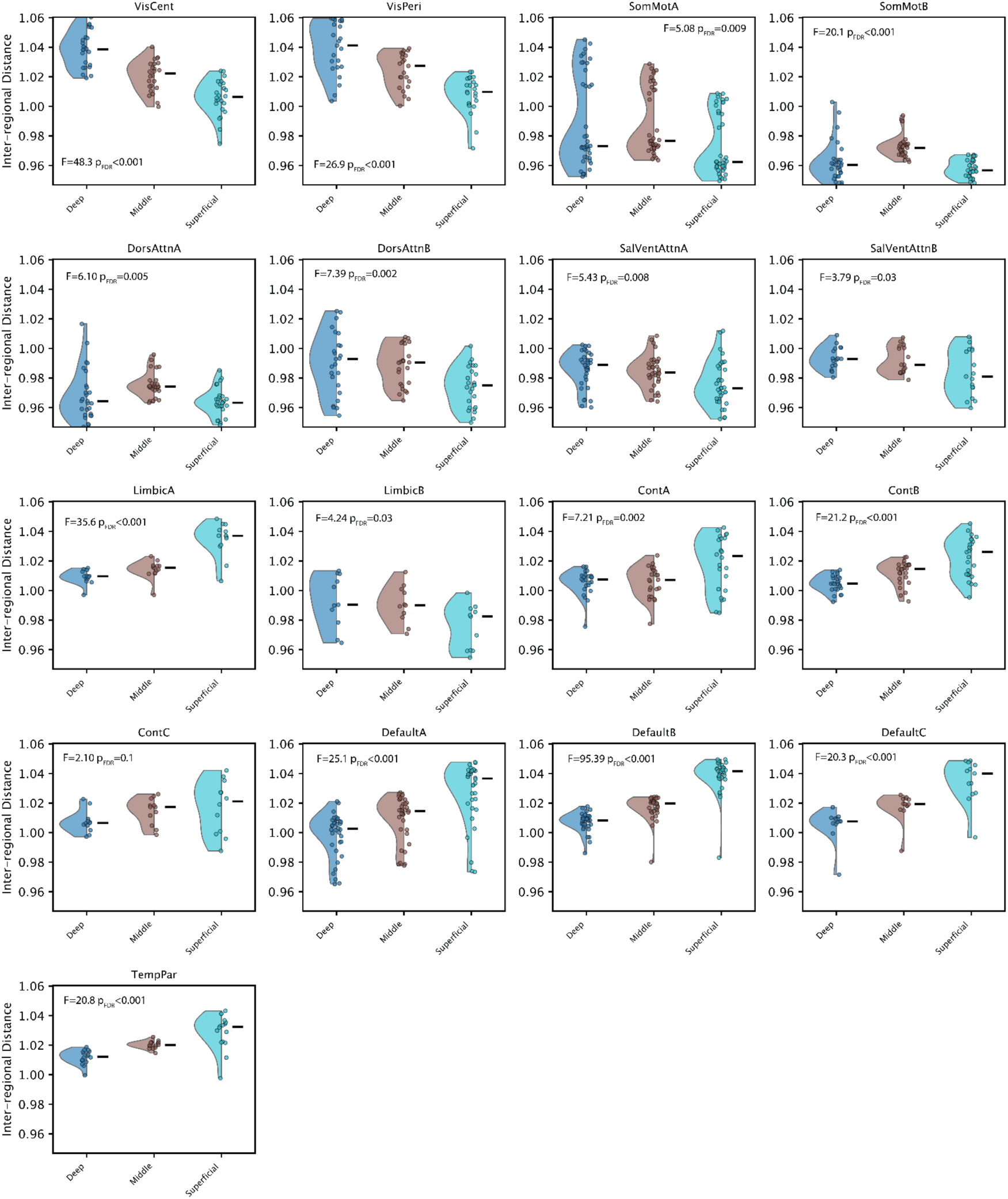
Yeo-17 network solution. Layer-specific inter-regional dissimilarity indices mapped per network. Each point represents a parcel within a given network. Statistics show tests for layer effects using a one-way ANOVA (FDR-corrected across networks).

**Supplementary Figure 2.**
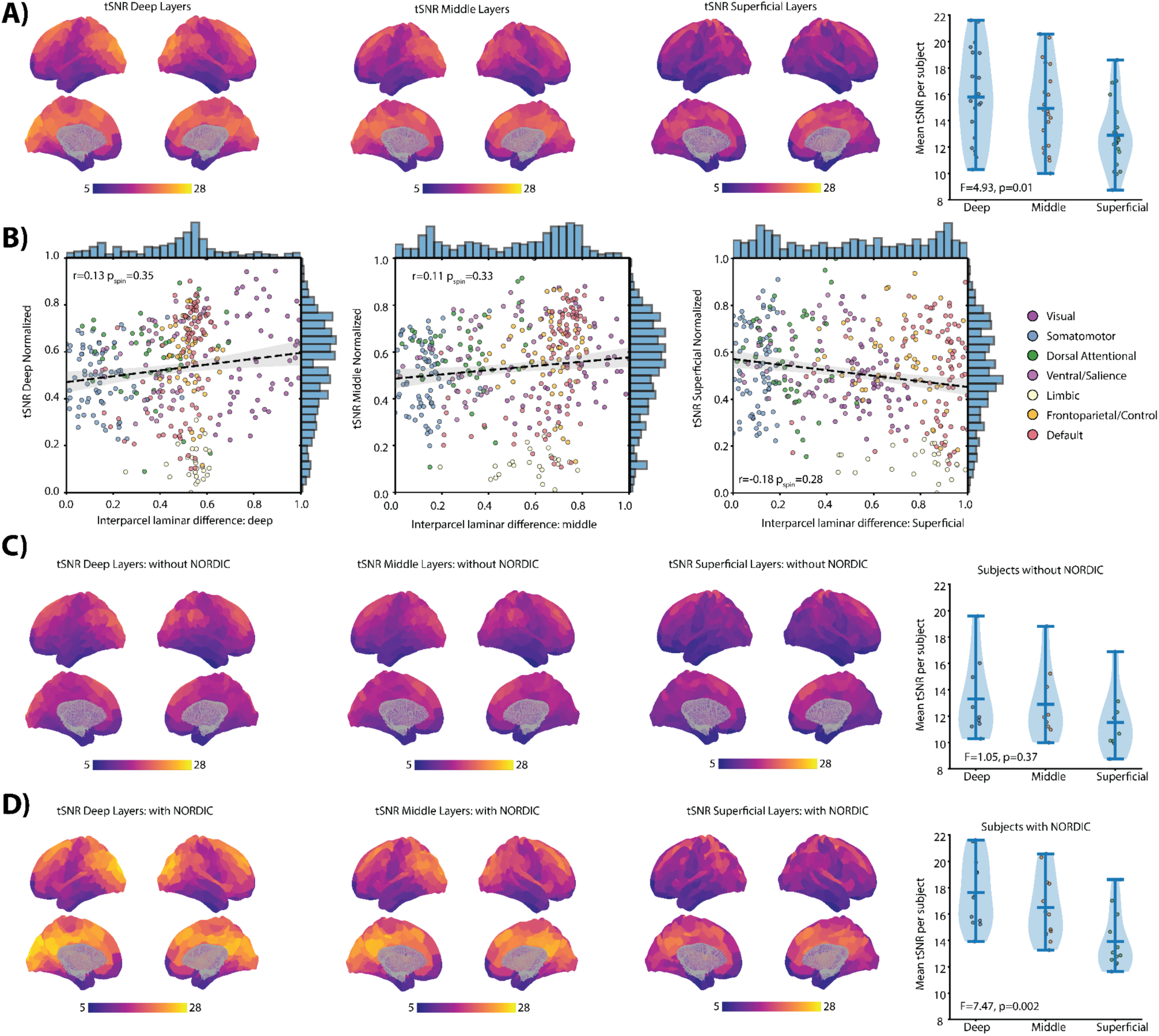
Temporal SNR. **A)** Maps: average tSNR across all subjects estimated per layer per Schaefer-400 parcel. Left: average tSNR across parcels per subject per layer. **B)** Correlation between tSNR of each layer and its corresponding inter-regional dissimilarity index. **C)** Same as A) but for subjects without NORDIC. **D)** Same as A) but for subjects with NORDIC.

**Supplementary Figure 3.**
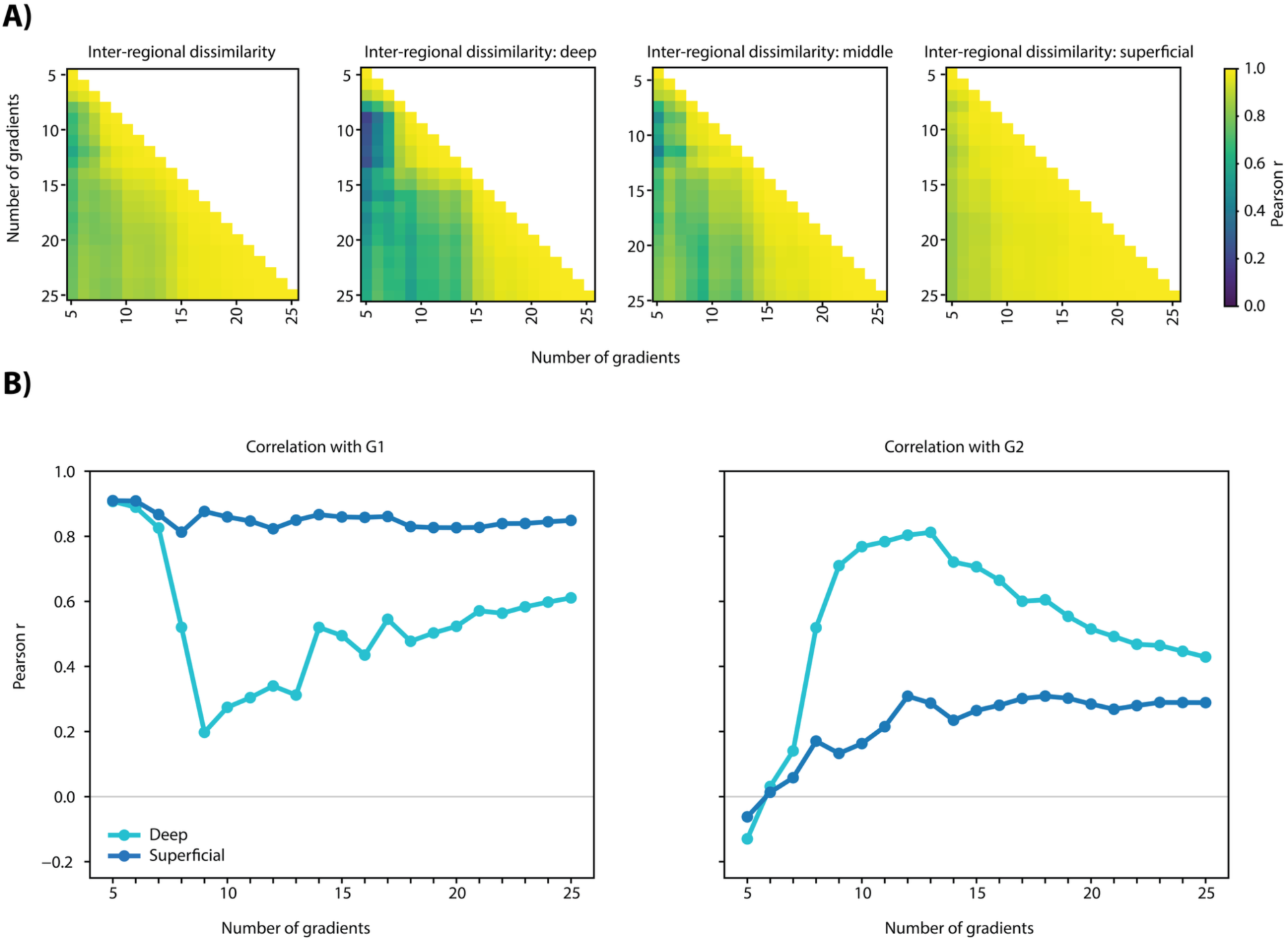
Dissimilarity index robustness. **A)** Correlations between dissimilarity indices calculated on the highest 5-25 gradients. **B)** Correlation of the superficial and deep indices calculated on the highest 5-25 gradients with G1 (left) and G2 (right).

**Supplementary Figure 4.**
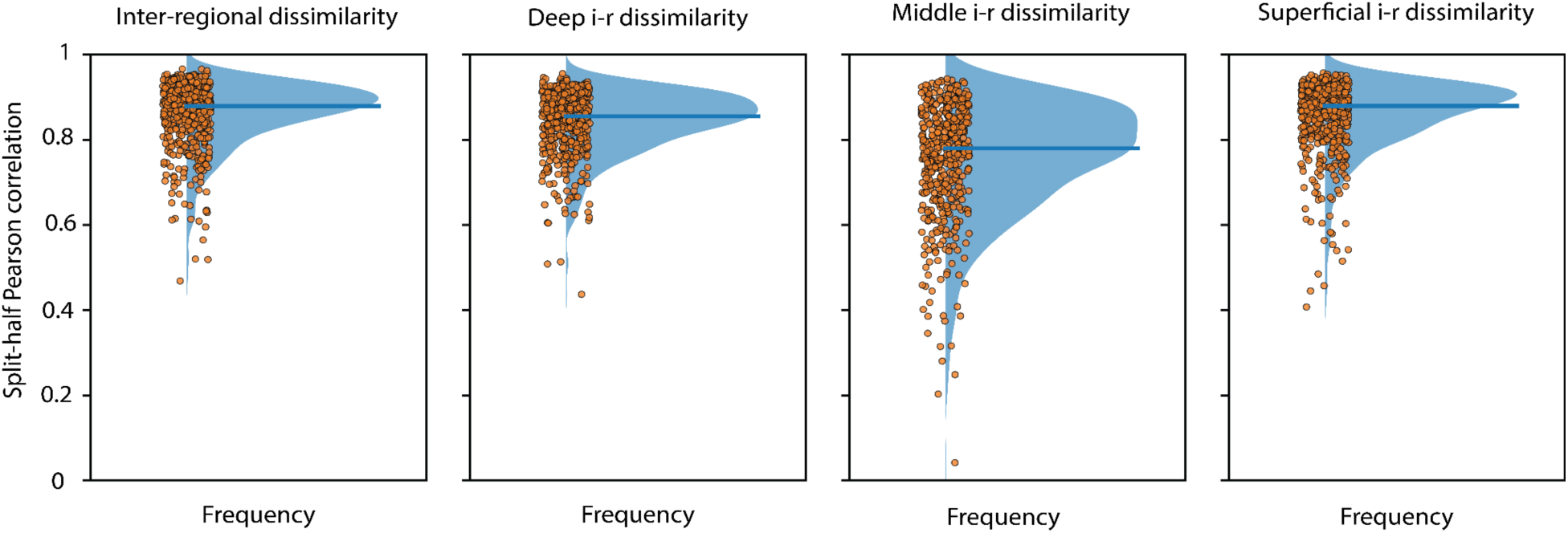
Split-half robustness. Split-half correlation of original cohort. Subjects were split in two groups and dissimilarities were computed for each group and correlated per pair; 500 iterations were performed. Points indicate a correlation value per iteration, while the curve shows the distribution with the horizontal line denoting the median. Inter-regional dissimilarity: μ = 0.88, σ = 0.077; deep dissimilarity: μ = 0.84, σ = 0.075; middle dissimilarity: μ = 0.76, σ = 0.13; superficial dissimilarity: μ = 0.85, σ = 0.086

**Supplementary Figure 5.**
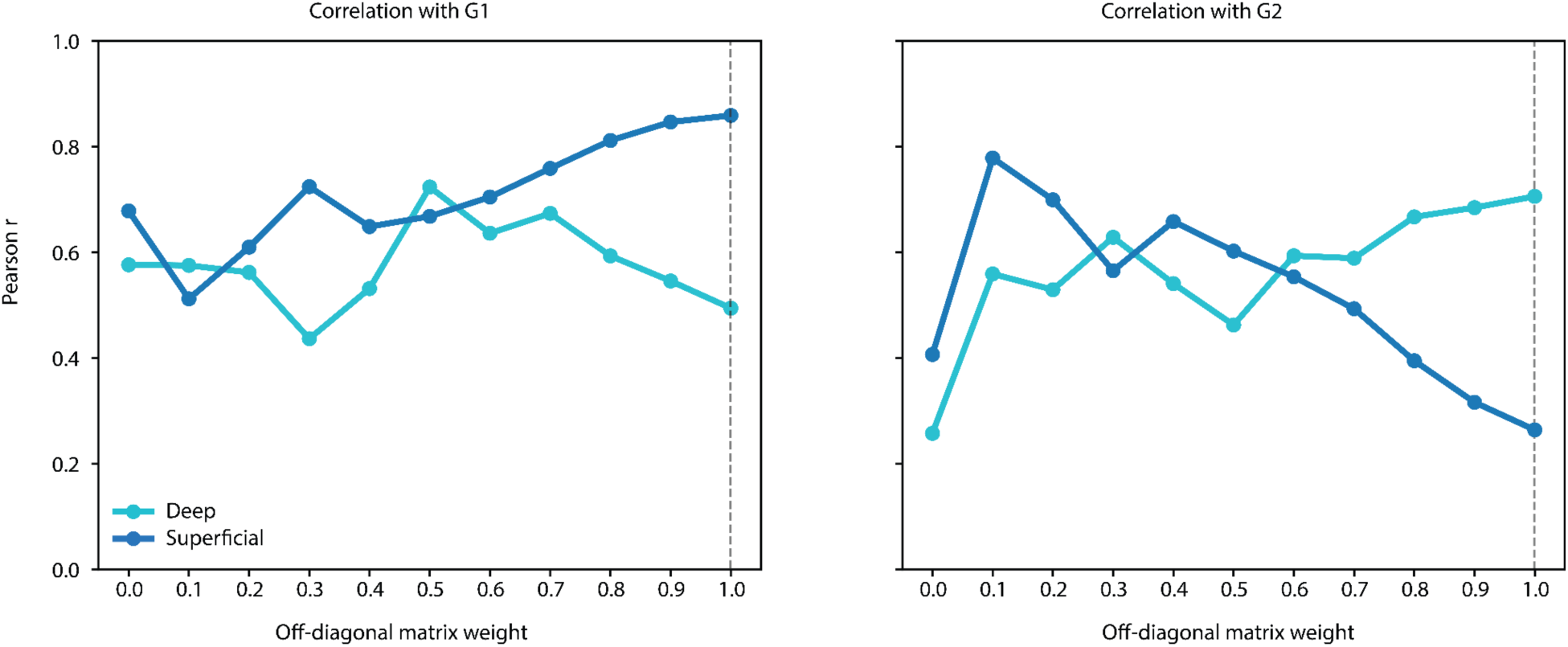
Identity matrix diagonal weight. Correlation of the superficial and deep indices with G1 (left) and G2 (right), as a function of the off-diagonal identity matrix’s diagonal weight. The original adjacency matrix had off-diagonals with a weight of 1, indicative of a strong coupling within each parcel across its layers.

**Supplementary Figure 6.**
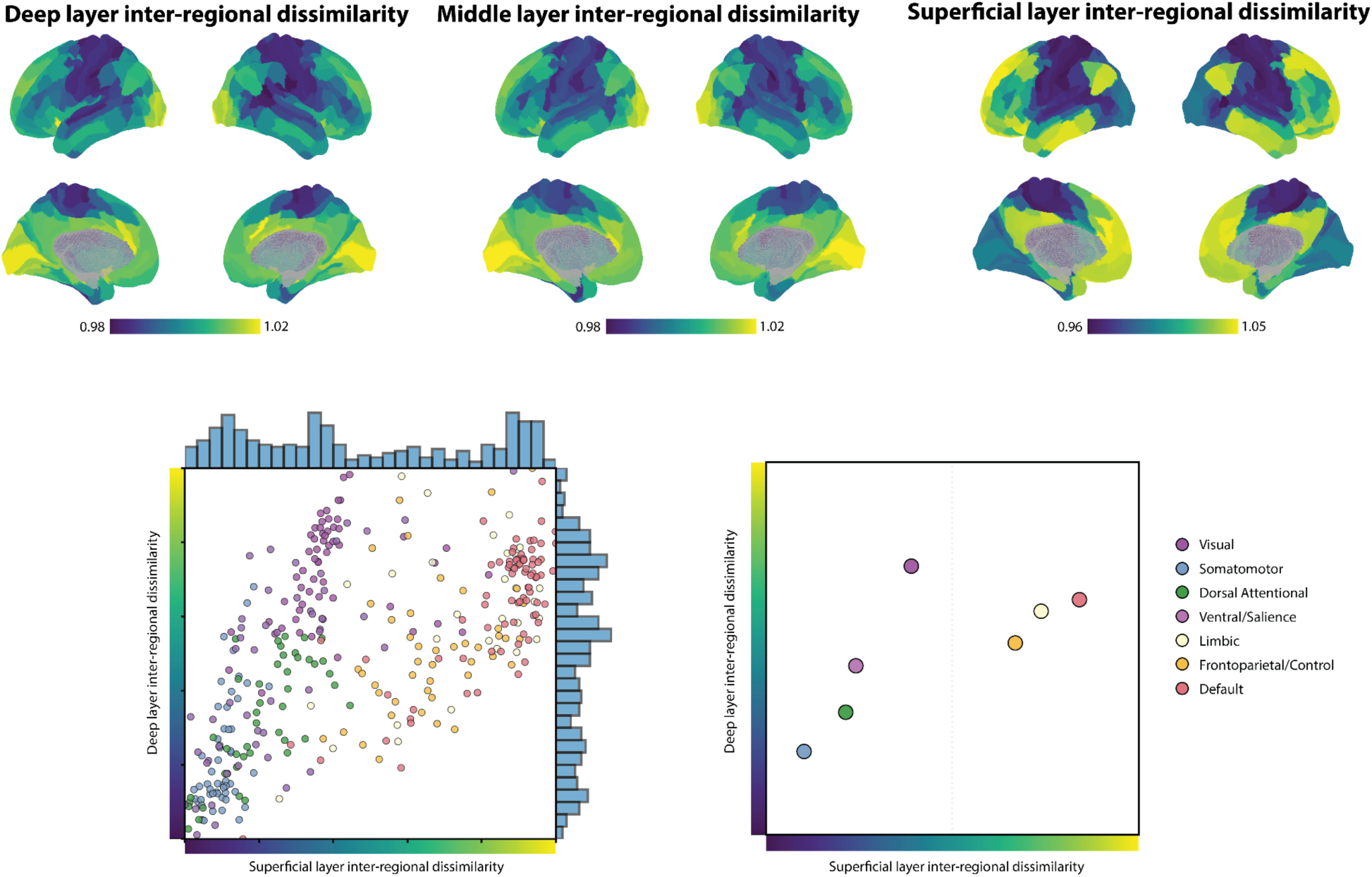
Layer-resolved resting-state connectivity and laminar dissimilarity indices in HCP MMP 1.0. Top: Layer-specific inter-regional dissimilarity computed on 15 gradients. Bottom left: Relationship between superficial and deep inter-regional indices across parcels. Each point is a parcel colored by its resting-state network using the seven network Yeo parcellation. Bottom right: Resting-state network centroids.

